# Soil Carbon Sequestration in *Nothofagus obliqua* Forests with Different Canopy Cover Levels under Silvopastoral Management

**DOI:** 10.1101/2025.01.11.632554

**Authors:** Camila Ramos, Erick Zagal, Salme Timmusk, Francis Dube, Leandro Paulino, Juan Ortiz, Jean Intriago, Juan Pablo Fuentes

## Abstract

Agroforestry contributes to slowing deforestation, favouring ecosystem regeneration and improving land use sustainability. In this study, we evaluated the effect of agroforestry on the recovery of degraded soils in a native forest and its capacity to capture and stabilize carbon (C) and nitrogen (N) in the soil. We conducted the study in a native forest of *Nothofagus obliqua* (oak) located in Ranchillo Alto (37°04′52″ S, 71°39′14″ W), in the Ñuble region, Chile. This site includes three silvopastoral systems with different levels of forest degradation: open (Op), semi-open (SOp) and semi-closed (SC), together with a control treatment without silvopastoral treatment (Ctr). The analysis was conducted at four soil depths (0-10, 10-20, 20-30 and 30-60 cm). To evaluate carbon and nitrogen sequestration and fixation in relation to the different forest cover treatments, we performed chemical, physical and biological soil analyses. We also included a physical fractionation by particle size to separate soil organic matter (SOM) into meaningful fractions with different characteristics and dynamics: particulate organic matter (POM) and organic matter associated with the mineral matrix (MAOM). Our results indicated that Op, despite being the most degraded condition, showed the highest values of C, N and non-oxidizable carbon (Cnox). This condition could be associated with the history of potato burning in this sector, which generates stable pyrogenic carbon in the soil. Analyses of C and N, as well as Cnox in the MAOM fraction, and C stock revealed that the three silvopastoral systems showed higher values than the control treatment without silvopastoral treatment. This suggests that the adoption of silvopastoral practices can improve soil quality and contribute to long-term carbon sequestration. These results support the application of sustainable practices to mitigate soil degradation and enhance carbon sequestration in degraded ecosystems.

## 1. Introduction

In Chile, decades of overuse of natural resources have led to 37.8% of the national territory showing soil degradation, in moderate to severe conditions, concentrated mainly in the central part of the country (Casanova *et al.,* 2013; Flores *et al.,* 2010). This degradation is mainly caused by the felling of more than 44% of native forests, affecting more than 8 million hectares (Mha). The genus *Nothofagus* has been one of the most impacted, decreasing its original area by more than 70%. This reduction is mainly due to anthropogenic activities, including the expansion of urban areas, croplands, and forest plantations, although most of the surface has been cleared for the growth of pastures for livestock use, reaching up to 3 Mha (Lara *et al.,* 2012).

Given this scenario of increasing soil and environmental degradation, various sustainable practices have been promoted in recent years to mitigate the impact of human activities. Among them, silvopastoral systems (SPS) have played a fundamental role. According to several studies (Yasin *et al.,* 2024; Nair *et al.,* 2021; Udawatta *et al.,* 2017), the integration of trees in production systems reduces erosion, improves soil fertility and quality, protects biodiversity, and contributes to the mitigation of climate change through carbon sequestration, being able to store between 1.8 and 6.1 Mg of Soil Organic Carbon (SOC) per year (Udawatta *et al.,* 2017; Nair et al., 2010).

Recent studies have highlighted the importance of including native forests and agroforestry systems in conservation and climate change mitigation policies (Yasin et al., 2024; Ortiz et al., 2023). Since 2019, the Intergovernmental Panel on Climate Change (IPCC, 2019) has underlined the crucial role of these systems in carbon capture and their potential to reduce greenhouse gas emissions, conserve biodiversity and protect soil (Cai and Aguilar, 2021; Kay *et al.,* 2019). FAO (FAO, 2018) as well called for collaboration between the agricultural and forestry sectors to expand agroforestry practices, defining agroforestry as a key climate solution due to its capacity to store large amounts of soil organic matter (SOM), which helps the provision of multiple ecosystem services (Smith *et al.,* 2015). According to Contla (2022), in agroecosystems with a marked presence of woody species, high amounts of SOM are produced, which translate into annual net increases in ecosystem C, of which around 60% of the C is in the form of SOC.

Soil organic matter can be defined as the complex mixture of different compounds from plants and soil microorganisms (Stevenson, 1994). The physical separation of SOM by particle size operationally into particulate organic matter (POM) and matrix-associated organic matter minerals (MAOM) is an approach that helps to understand the distribution of SOM and responses to environmental change (Cambardella and Elliott, 1992; Follett *et al.,* 2015; Cotrufo *et al.,* 2019; Witzgall *et al.,* 2021; Heckman *et al.,* 2022). It is proposed that POM and MAOM have different formation pathways and mean residence times in soil. It is postulated that POM (labile fraction of C) comes from the fragmentation or depolymerization of plant litter, while MAOM (stable fraction of C) comes from the transformations or modifications carried out by soil microorganisms and/or their exoenzymes, which grants it different degrees of physicochemical protection (Lavallee *et al.,* 2020).

Understanding the mechanisms of soil organic matter (SOM) formation and stabilization, as well as its sensitivity to disturbances and environmental changes, is fundamental for predicting its future dynamics (Cotrufo and Lavalle, 2022). Representing SOM pools based on the processes that regulate their formation and stabilization is crucial for developing sustainable management strategies. In this context, silvopastoral systems stand out for their ability to restore degraded soils by facilitating the incorporation and recycling of nutrients through SOM contributions. However, knowledge about the implementation of these systems in degraded native forests, particularly at different soil depths, and their capacity to stabilize soil organic carbon (SOC) in the long term remains limited. Therefore, the objective of this study was to evaluate the effect of agroforestry on the recovery of degraded soils in a native oak (*Nothofagus obliqua*) forest and its ability to capture and stabilize soil carbon and nitrogen.

## 2. Materials and Methods

### 2.1 Description of the Study Area

The study was carried out on a state-owned property in the Ñuble region of Chile (37°04′52″ S, 71°39′14″ W; 1200-2000 m a.s.l.) that covers a total area of approximately 635 ha (figure 1). This corresponds to a native forest (*Nothofagu*s *obliqua*) that has suffered significant soil degradation due to logging and prolonged overgrazing by local farmers. Most of the surrounding properties do not have the capacity to produce sufficient for livestock consumption throughout the year. This overgrazing has prevented the natural regeneration of trees in the more open areas, affecting the quality and density of the stands (Dube *et al.,* 2016). In 2015, silvopastoral systems were implemented in the northern and southern sectors of the forest, with the aim of recovering the ecosystem value of the native forest and promoting sustainable rural economic practices among the community. This study was carried out in the period between 2023-2024 and focused on silvopastoral systems located in the southern sector, which comprises approximately 10 hectares. In this area, three silvopastoral systems were established with different levels of soil degradation, determined by the degree of previous disturbance of the forest.

**Figure 1:**
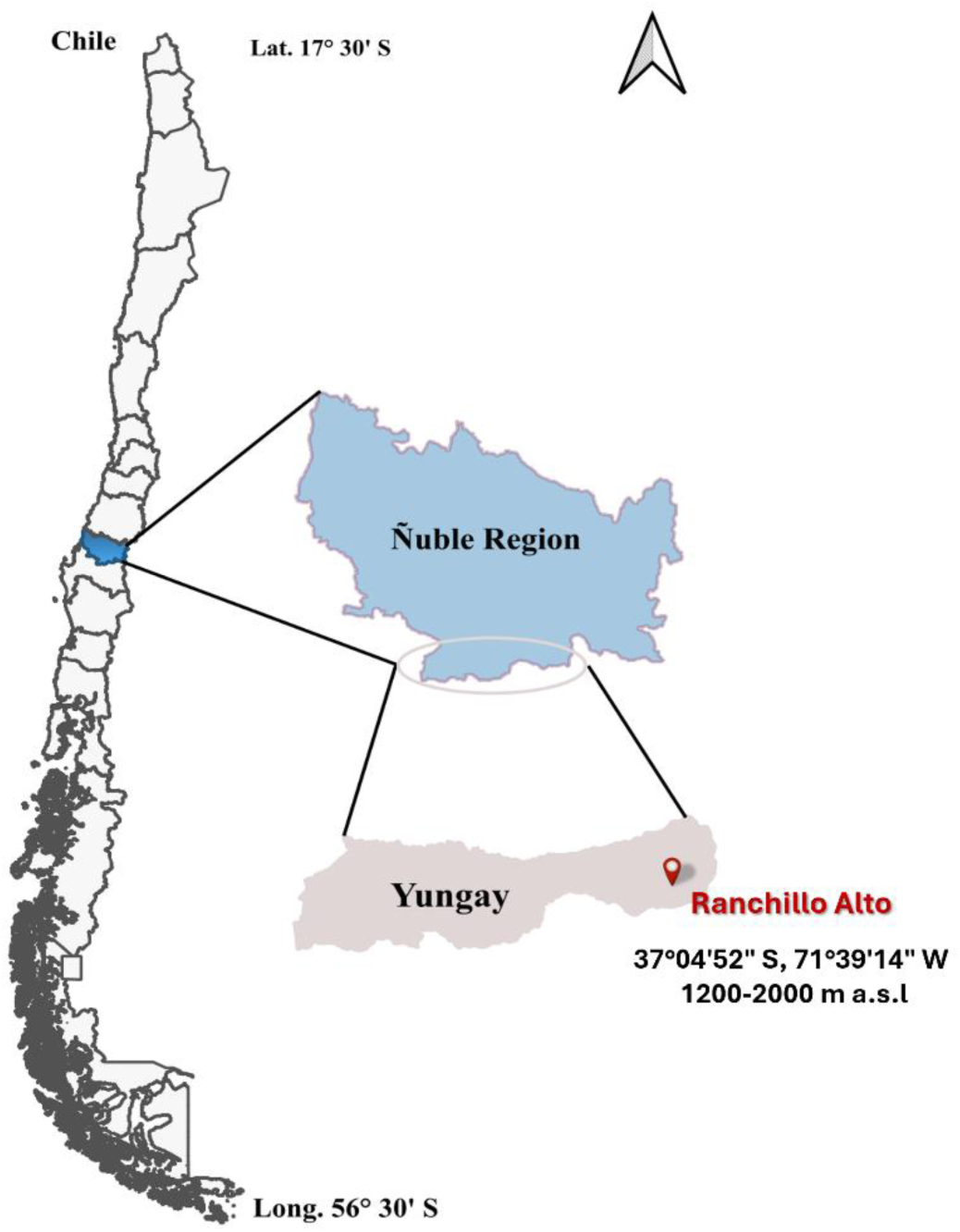
Map of the Ranchillo Alto native forest study site

### 2.2 Conditioning of experimental treatments

The effect of 4 conditions as experimental treatments was analysed. The sites will be classified according to the previous levels of canopy disturbance (+) as follows: open (Op) +++, semi-open (SOp) ++, semi-closed (SC) + and a degraded native forest control without silvopasture (Ctr) was also included (table 1).

**Table 1.** General information for each tree covers conditions in the silvopastoral systems located in the southern sector of Ranchillo Alto.

| Canopy opening | Ubicación | Number of plots y Superficie (ha) | Tree density (ha<sup>-1</sup>) | Forest species | Percentage of light | Previous degradation | Soil sampling |
| --- | --- | --- | --- | --- | --- | --- | --- |
| Ctr | 37°4'50" S, 72°26'30" W<br>1250 m.a.s.l | 3*1.33 |  | Roble (*Nothofagus obliqua*) |  | + | 4 depths (0-10, 10-20, 20-30 and 30-60cm) |
| OP | 37°14'51" S, 72°26'30" O<br>1250 m.a.s.l | 3*1.33 | 60 | Roble (*Nothofagus obliqua*) | 85-95% | +++ | 4 depths (0-10, 10-20, 20-30 and 30-60cm) |
| SOp | 37°14'50" S, 72°26'30" W<br>1250 m.a.s.l | 3*1.33 | 134 | Roble (*Nothofagus obliqua*) | 65-75% | ++ | 4 depths (0-10, 10-20, 20-30 and 30-60cm) |
| SC | 37°14'49" S, 72°26'30" W<br>1250 m.a.s.l | 3*1.33 | 258 | Roble (*Nothofagus obliqua*) | 45-55% | + | 4 depths (0-10, 10-20, 20-30 and 30-60cm) |

The native forest had the canopy openness conditions previously defined in 2015, (Ortiz et al., 2020), so we only validated the average percentages of solar radiation at ground level, as well as the leaf area indices to determine whether it was necessary to make minimal adjustments to tree density in case these levels have changed over time (see below).

To quantify and compare the percentages of cover for each treatment and their evolution over time, the results were corroborated with a second series of hemispherical photos taken using a Solariscope SOL 300B (Behling, Germany) (Annighöfer, *et al.,* 2015). These photos allowed the quantification of the percentages of cover for each treatment, according to the ranges of percentage of external light on the ground, which allowed determining that it was not necessary to remove new trees to obtain the desired luminosity levels in each treatment.

### 2.3 Soil sampling

A completely randomized design with three replicates (plots) randomly distributed at each tree cover level treatment was used for soil sampling.

Soil samples were taken in each plot, consisting of 5 random subsamples at depths of 0 - 10, 10 - 20, 20 - 30 and 30-60 cm, following the recommendations of Rumpel *et al.,* (2011). The samples were placed in a thermal box with ice to be transported from the forest to the laboratory and stored at 4°C. All samples were sifted with a 2 mm stainless steel sieve and then divided into 2 parts, one of which was air-dried for subsequent physical and chemical analyses and the other part was conditioned to 60% water filled pore space (WFPS humidity), ideal for biological analyses. All chemical, physical and biological analyses were carried out in the Spectroscopy Laboratory (Vis-IF) and Sustainable Soil Management of the Faculty of Agronomy of the University of Concepción.

### 2.4 Soil analysis

#### 2.4.1 Evaluation of physical parameters

For physical analyses, bulk density was determined by the cylinder method, true density by the pycnometer method, and texture was analysed by the Bouyoucos hydrometer method, all following the methodology proposed by Sandoval *et al*. (2012).

#### 2.4.2 Evaluation of chemical parameters

Chemical analyses of the soil, including pH, OM N, P, K Ke, Ca i, Al i, effective cation exchange capacity (CICE), S Mg, were performed according to the protocol of Sadsawka *et al*. (2006).

#### 2.4.3 Evaluation of biological parameters

Total enzymatic activity in soil was assessed by fluorescein diacetate (FDA) hydrolysis, a method that estimated the enzymatic activity of hydrolytic enzymes involved in organic matter degradation. For analysis, 1.0 g of moist soil was weighed into screw-capped test tubes (triplicates plus a blank). 9.9 mL of sodium phosphate buffer was added to the samples and 10 mL to the blank, followed by 0.1 mL of fluorescein diacetate (FDA) only to the samples. After vortexing, the tubes were incubated at 25°C for 1 hour. After incubation, they were cooled in an ice bath. For colorimetry, 10 mL of acetone was added to each tube, vortexed, and filtered. The absorbance of the filtrate was measured at 490 nm against a reagent blank prepared with acetone and distilled water. (Alef, 1995).

The activity of β-glucosidase, a key component of carbohydrate degradation in soil, was assessed using the method Tabatabai (1994), based on the release of p-nitrophenol after incubating the soil with p-nitrophenyl-β-D-glucopyranoside (25 mM) in MUB-HCl (pH 6) at 37°C for 1 hour. It was then cooled on ice, centrifuged at 6000 rpm for 5 minutes, and after the addition of 0.5 M CaCl₂ and THAM-NaOH buffer (pH 12), the absorbance at 400 nm was measured. Both the absorbance of β-glucosidase and FDA enzymatic activity were measured using a UV-visible spectrophotometer (AA3, BRAN + LUEBBE, Norderstedt, Germany).

For soil respiration, 20 g of moist soil conditioned at 60% WFPS per treatment (in triplicate) was weighed into a Falcon tube. These vials were sealed with specialized caps that allow gas (CO2) extraction using a precision syringe, which was then injected into a LICOR gas analyzer to perform microbial respiration analysis. The vials with soils were kept in an incubation chamber at 22°C for 14 days. This requires that the soil has been conditioned at 60% WPFS. Basal soil CO2 emission is calculated at 3, 5, 7, 10 and 14 days of incubation. To perform the gas extraction process, a homogenization of the Falcon’s headspace was previously performed by extracting 1 ml of gas and injecting it again three times. The 1 mL gas sample was then injected into a unidirectional dual-wavelength non-dispersive infrared (NDIR) gas analyzer (LI-820 CO2 Analyzer), using a modification as described by Craine *et al*. (2010).

#### 2.4.4 Soil physical fractionation

The physical fractionation of the soil was conducted following the method described by Lavallee et al. (2019) to enhance understanding of the stable carbon fraction. Briefly, as outlined by Lavallee et al. (2019), soils were sieved to 2 mm, and 5 g of oven-dried soil were shaken with 15 mL of 0.5% sodium hexametaphosphate solution and five 1-mm glass beads for 18 hours to disperse the soil. The dispersed soil was then rinsed through a 53 µm sieve, with the fraction passing through (<53 µm) collected as MAOM, while the remaining material was classified as POM.

#### 2.4.5 Total Carbon, Oxidizable Carbon, Non-Oxidizable Carbon, and C stocks calculations

Total carbon (TC) and total nitrogen (N) were determined in each of the soil samples and their fractions using the dry combustion method based on the Dumas principle (Wright et al., 2001). The different samples and their fractions were subjected to chemical oxidation with sodium dichromate dihydrate to determine the proportion of oxidizable carbon (Cox), following the Walkley-Black method (Sadsawka *et al*. 2006). Since there is no inorganic carbon in the volcanic soil, TC = total organic carbon (OC). Non-oxidizable organic carbon (Cnox) was determined by the difference between total carbon and non-oxidizable carbon (Cnox = TOC - Cox).

The carbon stock (C stock) for each soil layer was calculated by multiplying the organic carbon concentration (% OC) by the bulk density (BD, g cm⁻³) and the thickness of the sampled soil layer (cm). The formula used was:

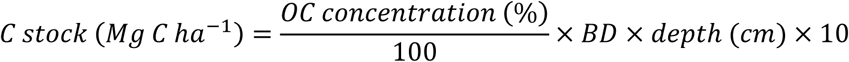

### 2.5 Statistical analysis

To evaluate the effect of open (Op), partially open (SOp), partially closed (SC) and Control (Ctr) canopy cover on the different chemical, physical and biological analyses at the different depths (0-10; 10-20; 20-30 and 30-60 cm), a two-way ANOVA was performed. And for significant differences, Tukey Post Hoc analyses were performed, using the R program (R Core Team 2021). A principal component analysis (PCA) was performed to identify patterns in the variability of the variables studied and to reduce the dimensionality of the data. This analysis allowed observing the distribution of the samples based on the treatments and soil depths, and to evaluate the relationship between the different soil variables studied. The PCA was performed using the R program (R Core Team, 2021), employing the prcomp function with previously centered and scaled data.

## 3. Results and Discussion

Our physical analyses of the soil revealed that the apparent and true density, as well as the texture, remained within the typical range for volcanic soils under the different site conditions (Table 2) (Casanova et al., 2013).

**Table 2.** Physical Analysis Results.

| Condition/Depths | BD (g cm<sup>-3</sup>) | PD (g cm<sup>-3</sup>) | Sand (%) | Silt (%) | Clay (%) |
| --- | --- | --- | --- | --- | --- |
| Ctr (0-30) | 0.66±0.02Aa | 2.26±0.00Aa | 34.63±1.10Aa | 42.57±0.40Ab | 22.79±0.77Aa |
| Ctr (30-60) | 0.63±0.02Ab | 2.24±0.01Ab | 36.60±4.09Aa | 46.13±3.77Aa | 17.27±0.40Ab |
| OP (0-30) | 0.61±0.02Aa | 2.14±0.02ABb | 37.17±0.37Aa | 45.80±0.37Ab | 17.03±0.68Ba |
| OP (30-60) | 0.76±0.08Ab | 2.21±0.00ABa | 33.80±2.36Aa | 48.87±0.37Aa | 17.33±1.41Bb |
| SOp (0-30) | 0.61±0.03Aa | 2.09±0.01Bb | 40.57±1.23Aa | 43.90±1.33Ab | 15.50±0.99Ba |
| SOp (30-60) | 0.72±0.06Ab | 2.12±0.09Ba | 35.70±1.04Aa | 50.03±1.59Aa | 14.27±0.37Bb |
| SC (0-30) | 0.60±0.01Aa | 2.06±0.01Bb | 39.77±3.01Aa | 43.03±0.82Ab | 17.20±2.12ABa |
| SC (30-60) | 0.64±0.06Ab | 2.20±0.05Ba | 37.13±2.53Aa | 45.07±1.17Aa | 17.80±2.47ABb |

n :24 and p < 0.05. Different capital letters mean significant differences between conditions, while lowercase letters indicate significant differences between depths. BD: apparent density, PD: real density, % Sand: percentage of sand, % Silt: percentage of silt and % Clay: percentage of clay

Individually, the apparent density showed representative values for Andisols ranging between 0.60 and 0.76 g/cm³. The density variation depended more on the soil depth than on the canopy opening, being more significant in the surface horizon. These observations coincide with the results of Gómez *et al*. (2022), who reported similar findings in Andisols soils from forests under silvopastoral management in Argentine Patagonia.

In addition, the real density, like the apparent density, presented variations in terms of soil depth, giving significant differences. The actual density ranged from 2.06 to 2.21 g cm ^-3^ with a mean value of 2.1 g cm ^-3^, which is within representative ranges of volcanic soils rich in OM, according to Nissen *et al*. (2005) and Ortiz et al. (2020) who estimated ranges of 1.9 to 2.1 (0 to 15 cm) and 1.9 to 2.0 (0 to 20 cm) for PD in forest soils (0–15 cm).

The soil texture, like the rest of the physical variables, did not show differences in terms of canopy opening levels, but did show significant differences between soil depths for sand and silt. The results found for texture coincide with those reported by Gómez *et al*. (2022) Chemical analyses of the soil generally showed results representative of Andisol soils (Table 3). The observed pH values are slightly acidic, fluctuating between 5.41 and 5.66, a typical characteristic of Andisol soils (Arifin *et al.,* 2022). Lower pH values were recorded in the open system at both depths, which could be attributed to higher acidity stimulated by a high OM content (Nanzyo 2002).

**Table 3.** Chemical Analysis Results.

|  | pH (H2O) | OM<br>(%) | N<br>(mg kg<sup>-1</sup>) | P<br>(mg kg<sup>-1</sup>) | K<br>(mg kg<sup>-1</sup>) | CICE<br>(cmol kg<sup>-1</sup>) | Al i<br>(cmol kg<sup>-1</sup>) | Ca<br>(%) | Mg<br>(%) |
| --- | --- | --- | --- | --- | --- | --- | --- | --- | --- |
| **Ctr (0-30)** | 5,45±0.05 | 14,51 | 40,77±4.88 | 1,88±17.76 | 110,28±0.04 | 3,96±1.52 | 0,09±0.01 | 3,01±1.34 | 13,75±2.04 |
| **Ctr (30-60)** | 5,66±0.04 | 9,07 | 8,30±3.18 | 1,23±8.03 | 37,60±0.02 | 1,37±0.72 | 0,03±0.01 | 0,95±0.59 | 17,24±2.29 |
| **OP (0-30)** | 5,45±0.05 | 22,07 | 21,10±0.05 | 0,67±0.05 | 91,17±0.05 | 2,14±0.05 | 0,18±0.05 | 1,43±0.05 | 12,85±0.05 |
| **OP (30-60)** | 5,41±0.04 | 14,38 | 14,40±0.04 | 0,53±0.04 | 45,27±0.04 | 0,86±0.04 | 0,07±0.04 | 0,49±0.04 | 17,81±0.04 |
| **SOp (0-30)** | 5,62±0.21 | 17,49 | 20,57±0.09 | 0,73±19.36 | 62,57±19.36 | 2,36±1.57 | 0,07±0.04 | 1,83±1.43 | 12,89±2.77 |
| **SOp (30-60)** | 5,63±0.08 | 10,52 | 10,83±0.09 | 0,67±22.66 | 46,60±22.66 | 1,20±0.56 | 0,01±0.00 | 0,83±0.48 | 15,82±4.46 |
| **SC (0-30)** | 5,50±0.18 | 18,37 | 26,97±0.14 | 0,90±21.32 | 90,50±21.32 | 3,43±1.52 | 0,14±0.09 | 2,62±1.43 | 12,30±1.63 |
| **SC (30-60)** | 5,44±0.12 | 12,67 | 17,30±0.05 | 0,57±38.09 | 81,87±38.06 | 1,92±0.84 | 0,09±0.05 | 1,29±0.69 | 14,54±2.89 |

OM showed a decrease with depth in all treatments, an expected pattern given that OM is mostly concentrated in the surface horizon due to the accumulation of leaf litter and roots. In the surface horizon (0-30 cm), the OP system presents the highest OM value (22.07%), followed by SC, SOp and Ctr respectively, reflecting the direct contribution of organic matter in the silvopastoral treatments compared to the control. The OM values are relatively higher than those reported in other studies (Gomez *et al.,* 2022), this is mainly due to the location of the forest in the foothills, where low temperatures throughout the year slow down the decomposition processes of soil microorganisms (Doetterl *et al.,* 2015).

Available N decreased with depth in all treatments, which is consistent with that presented in organic matter. In silvopastoral systems, nitrogen levels were considerably higher than in the control (Ctr), especially in the surface horizon. These results suggest that silvopastoral systems support microbial activity and nitrogen mineralization, which increases the availability of this nutrient for plants. This is consistent with previous studies indicating that this type of practice can improve nitrogen retention and availability in soils, mainly due to the incorporation of organic waste from animals in these systems, which provides an additional source of nitrogen (Renwick *et al.,* 2025). P presented low values in all treatments, this is a common characteristic in Andisols, due to the high phosphate adsorption capacity of amorphous minerals present in these soils (Redel *et al.,* 2008).

The CICE values were highest at the depth of 0-30 cm. At this depth, Ctr had the highest value, with 3.96 cmol/kg, followed by SC, with 3.43 cmol/kg. This difference can be attributed to the fact that these two conditions are the least disturbed, according to the previous levels of canopy degradation (+). This result is closely related to the values observed for K, Ca and Mg, since the CICE favours the accumulation of amorphous minerals, which contribute significantly to the cation retention capacity (Dahlgren & Saigusa, 2018). Likewise, this capacity is directly associated with the availability of essential cations for plant nutrition, whose presence can be influenced by the mineralogical composition, particularly by the higher clay content found in Ctr and SC.

FDA activity is a measure of overall enzymatic activity in soil, used as an indicator of total microbial activity. Figure 2 shows the results of microbial biomass activity as a function of canopy openness and soil depth. FDA hydrolysis analyses reveal significant differences both between different soil depths (ANOVA F = 8.56; df = 3,32; p = 0.0003) and between different canopy openness conditions (ANOVA F = 22.91; df = 3,32; p = 0.0001). In addition, significant interactions were found (ANOVA F = 4.85; df = 9,32; p = 0.0004) for depth and canopy openness (Table S1).

**Figure 2:**
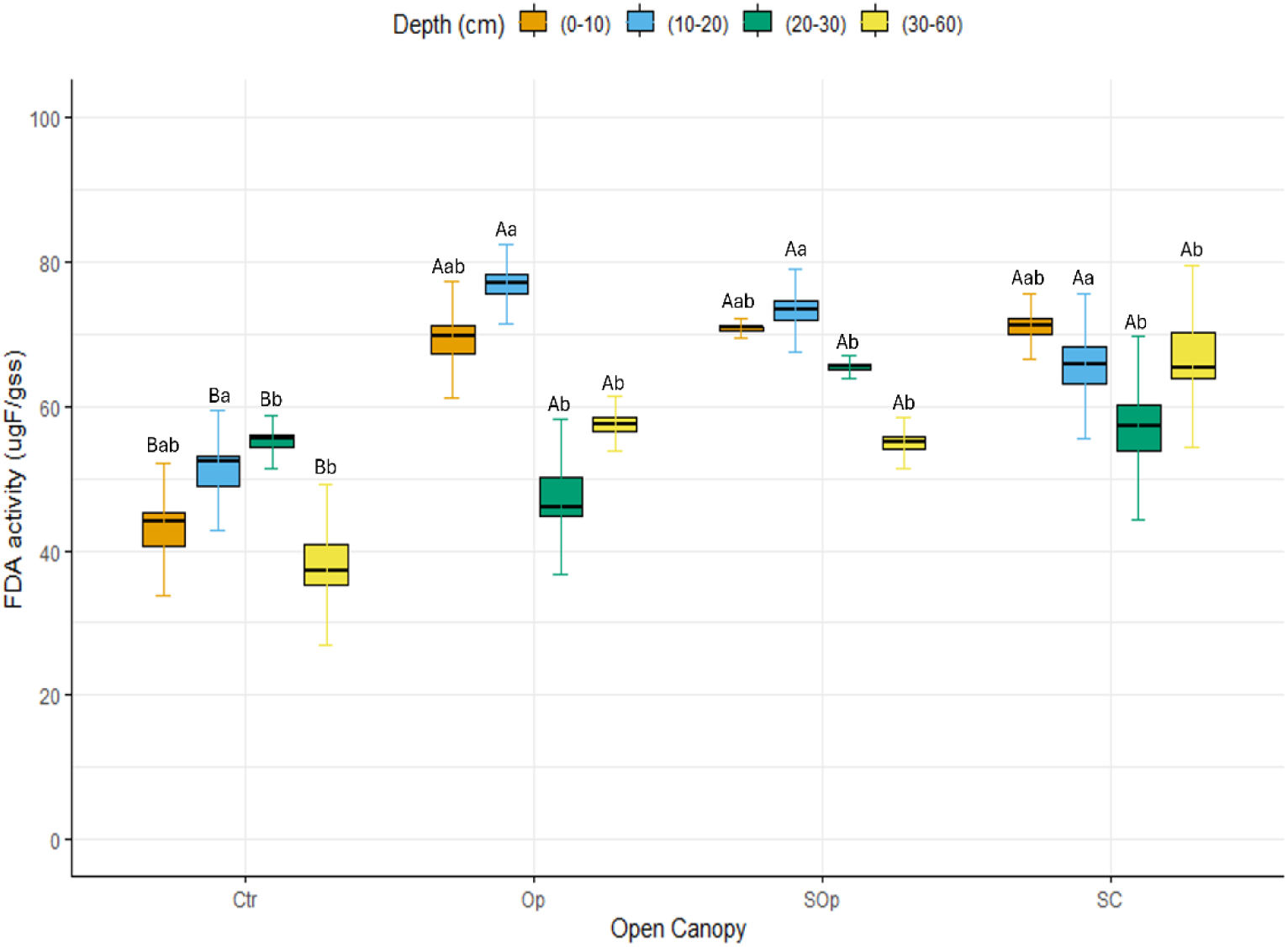
Microbial biomass activity determined by FDA hydrolysis as a function of canopy openness (Ctr: control; Op: open; SOp: semi-open; SC: semi-closed) and soil depth. Uppercase letters indicate significant differences between canopy conditions, while lowercase letters indicate significant differences between soil depths. Differences were considered significant according to Tukey’s test (p < 0.05).

It is observed that FDA activity decreases as soil depth increases, regardless of the type of management. This is related to the fact that soil microorganisms, which are responsible for the hydrolysis of FDA, are more concentrated in the surface layers of the soil, where there is greater availability of organic matter and nutrients. On the other hand, significant differences are observed between the plots with silvopasture compared to the control, which may suggest that including SPS helps soils recover their functional capacity after having been degraded.

The results obtained for the FDA in this study are consistent with those of previous research carried out at the site by Ortiz *et al*. (2023). However, they are higher than those reported by Reyes *et al*. (2011), who evaluated the biological activities of the soil in a relict forest in south-central Chile within a mixed forest community. Their results are very similar to those of the Ctr, while silvopastoral systems present higher values. This indicates that these systems promote greater biological activity in the soil, possibly due to the combination of plant species and management practices that favor the availability of organic matter and microbial diversity.

Like FDA activity, soil respiration showed significant differences between conditions (ANOVA F = 14.73; df = 3.32; p = 0.0001) and soil depths (ANOVA F = 106.64; df = 3.32; p = 0.0001) (Figure 3). The accumulated CO₂ fluxes, measured in a closed system for 14 days, were higher at shallow depths (0-10 cm) compared to the other depths for the four treatments. The SOp and SC conditions were those that showed the greatest increase in CO₂ emissions. This relationship could be influenced by variations in litter contributions, since a higher tree density leads to a greater incorporation of organic matter into the soil and a greater metabolic activity of microorganisms. In addition, closed systems limit the incidence of sunlight, which helps to preserve the humidity and temperature conditions in the litter, favouring the subsequent proliferation of fungi and bacteria.

**Figure 3:**
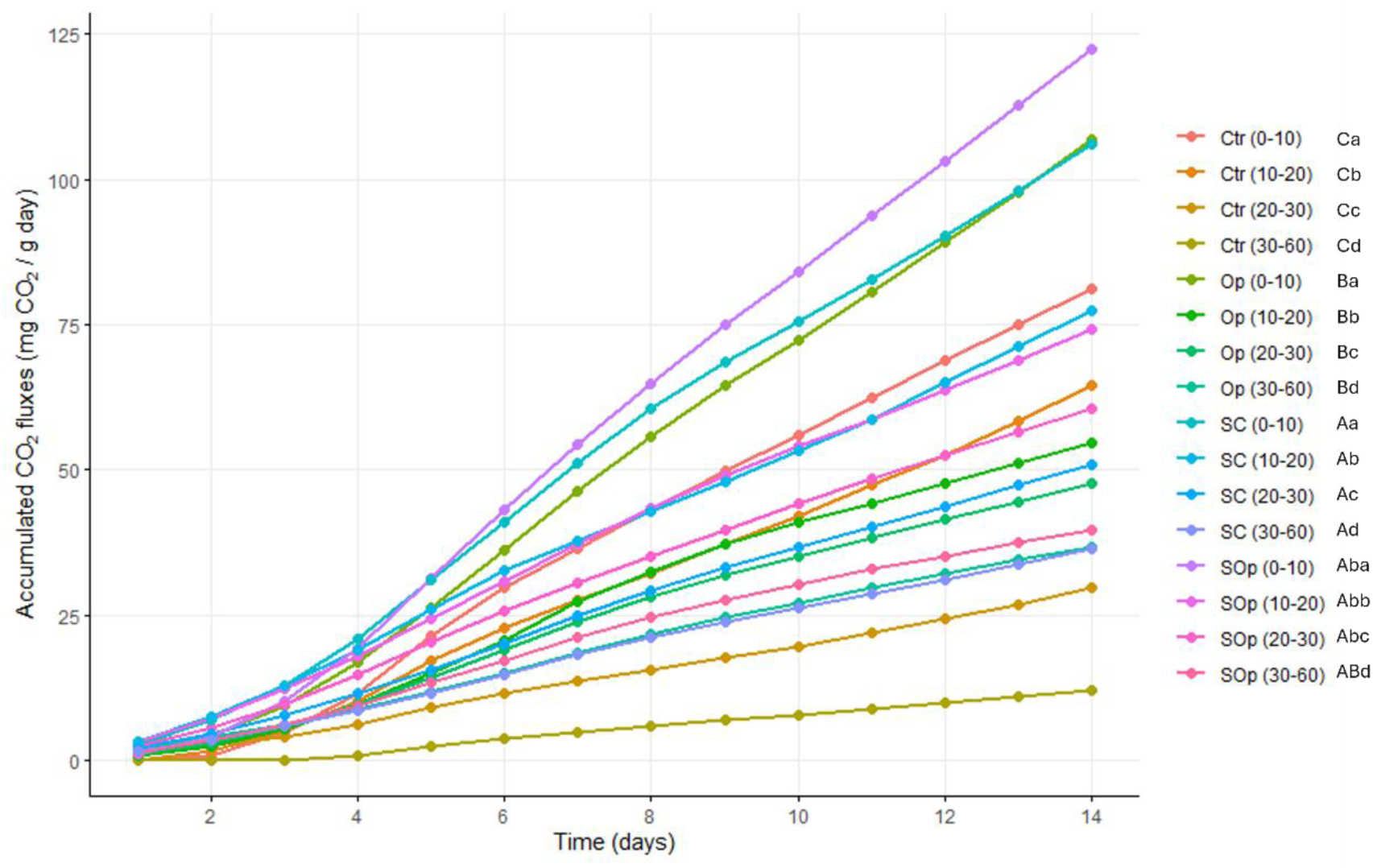
Cumulative soil respiration fluxes over a 14-day period. Each line represents a time series of CO2 measurements for different soil depths and treatments. Analysis of variance (ANOVA) was used to determine significant differences. Uppercase letters indicate significant differences between canopy conditions, while lowercase letters indicate significant differences between soil depths. Differences were considered significant at p < 0.05.

β-Glucosidase, unlike FDA activity and respiration, did not show significant differences between conditions (ANOVA F = 1.43; df = 3.32; p = 0.2521) (Figure 4). However, it was observed that OP˃ SC˃ SOp˃ Ctr showed a similar trend to the rest of the biological analyses. This is because this indicator is less sensitive to changes in soil conditions, since its activity is more related to specific processes of the carbon cycle, while FDA and respiration reflect a broader range of microbial metabolic activities and responses to environmental variations (Reyes *et al.,* 2011). Regarding depth, it did show significant differences, finding greater β-Glucosidase activity in the surface horizons. Since β- Glucosidase is an important indicator of the decomposition of organic matter, especially of polysaccharides such as cellulose, this result could have been conditioned by the fact that in the superficial horizons (0-10 cm) there is the greatest availability of carbon-rich substrates, such as those from the decomposition of leaf litter, and as one goes deeper into the profile (30-60 cm) the contributions of fresh organic matter and oxygen conditions decrease, which limits enzymatic activity.

**Figure 4:**
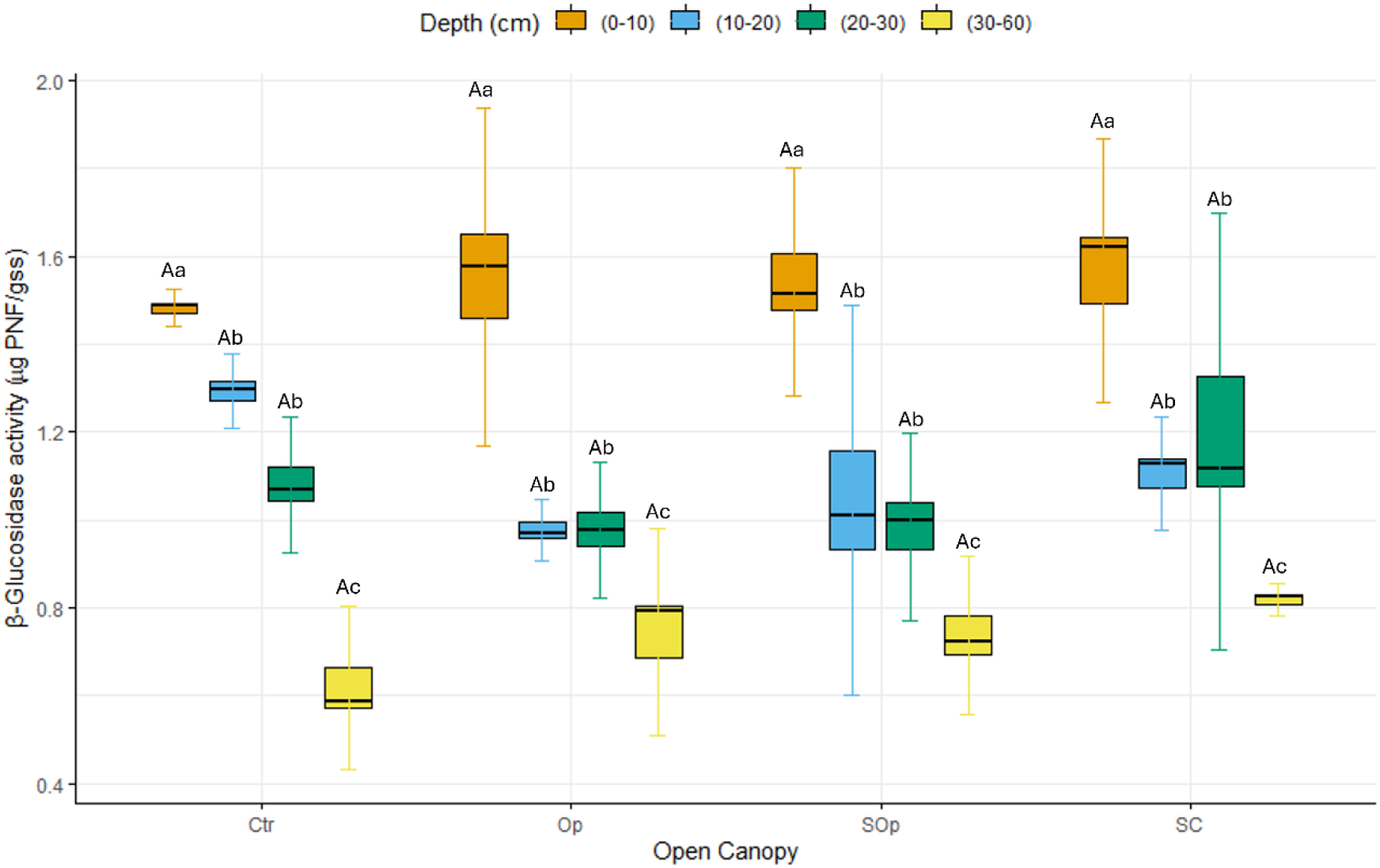
β-Glucosidase activity as a function of canopy openness and soil depth. Bars represent the mean of each variable, and whiskers indicate the standard error of the mean. Different capital letters mean significant differences between conditions, while lowercase letters indicate significant differences between depths. Analysis of variance (ANOVA) was used to determine significant differences. Differences were considered significant at p < 0.05.

The organic carbon content in the soil showed significant differences between the evaluated depths (ANOVA F = 25.93; df = 3.32; p = 0.0001) (Figure 5a). The highest concentrations were found in the surface horizons (0-10 cm). This pattern is consistent with the dynamics of carbon in soils, since the surface horizons receive greater contributions of leaf litter and other organic residues that, when decomposed, favor the accumulation of C and as the depth of the profile increases, the percentage of C decreases due to the lower amount of fresh organic matter and a more limited microbial activity.

**Figure 5:**
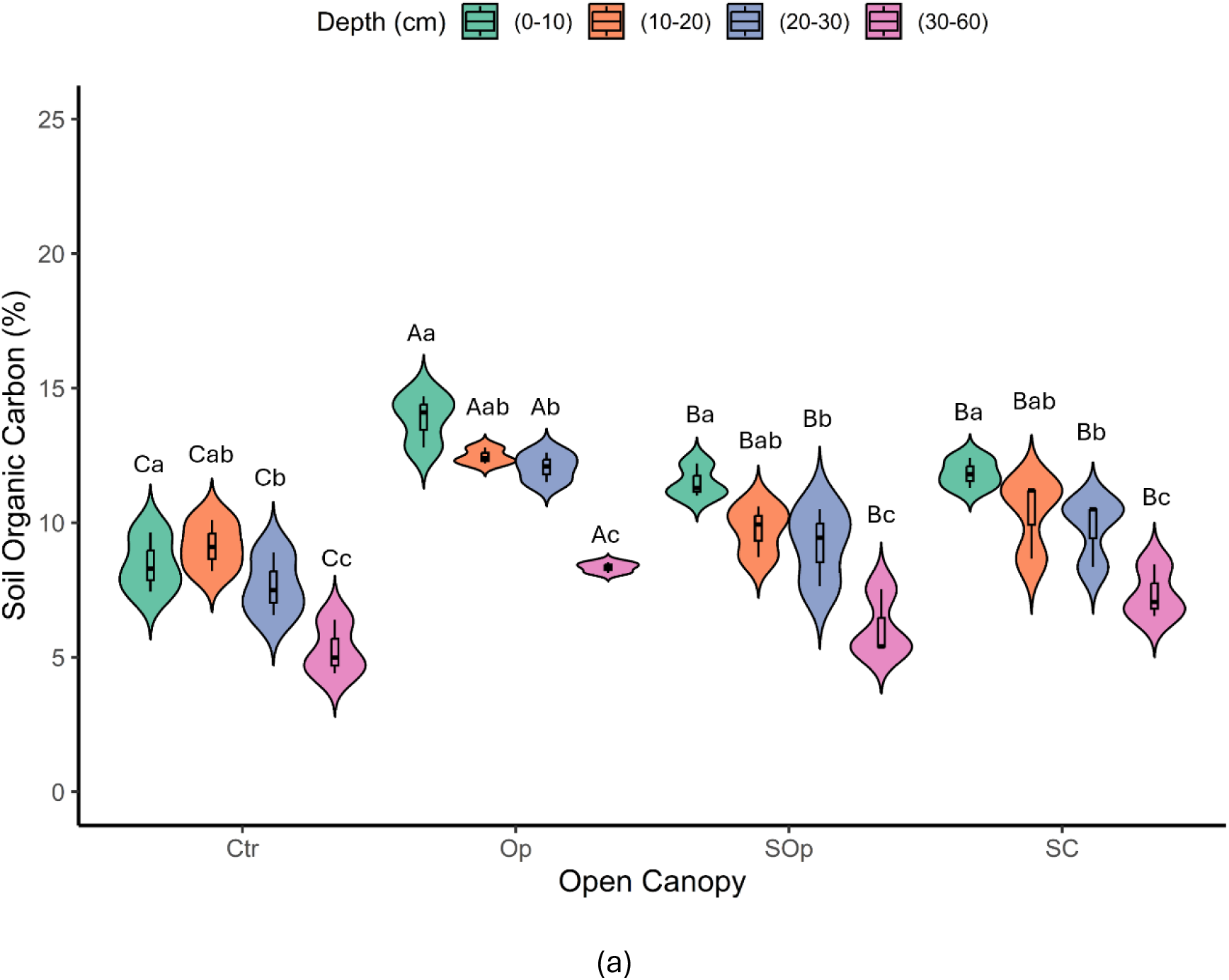
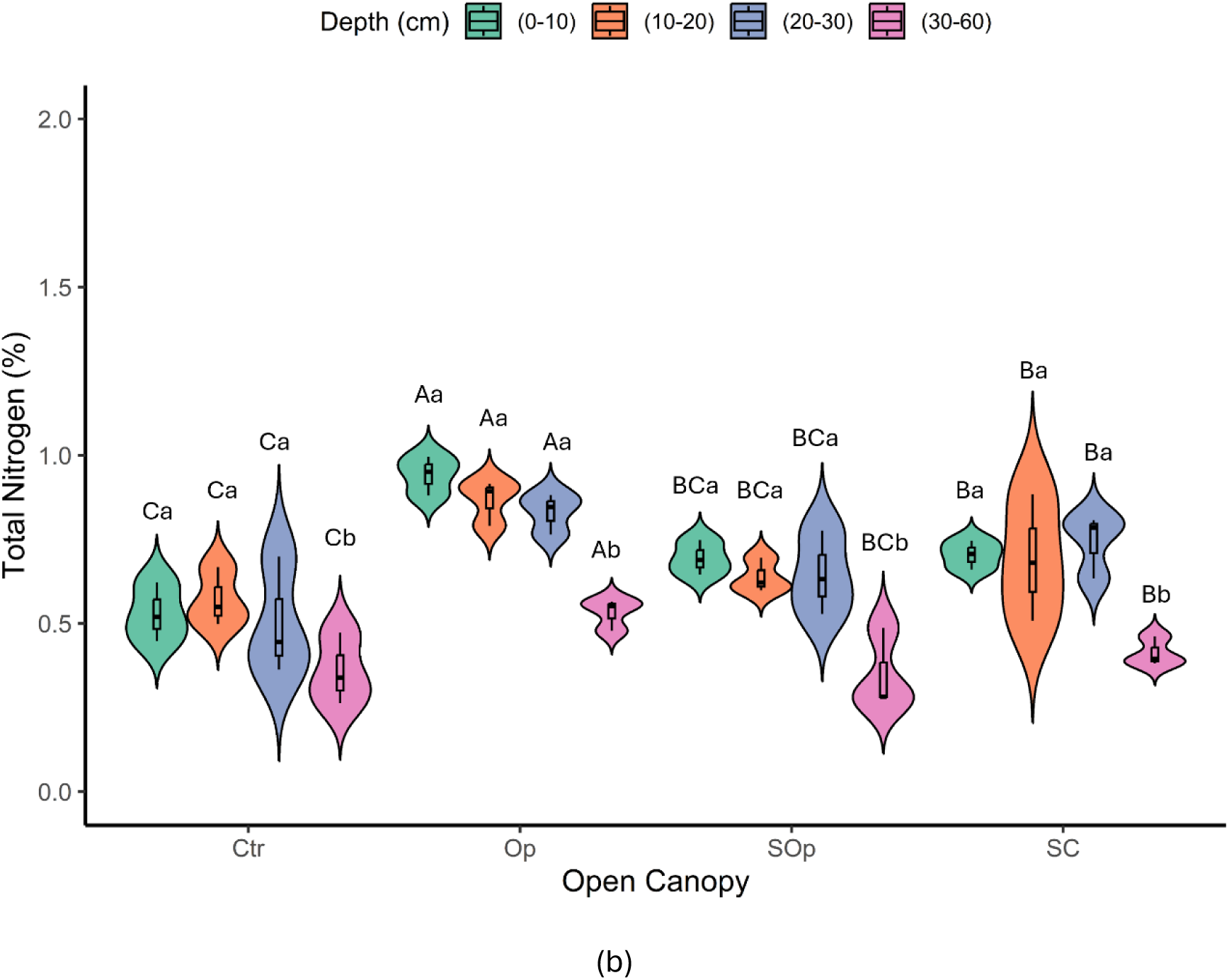
Violin plot of variables (a) Organic Carbon (%) and (b) Total Nitrogen as a function of factors canopy openness and soil depth. Bars represent the mean of each variable, and whiskers indicate the standard error of the mean. Different capital letters mean significant differences between conditions, while lowercase letters indicate significant differences between depths. Analysis of variance (ANOVA) was used to determine significant differences. Differences were considered significant at a p < 0.05 level.

On the other hand, significant differences were observed between the different canopy openings (ANOVA F = 34.38; df = 3.32; p = 0.0001). The Op canopy showed the highest concentrations of C, despite having the lowest tree cover and being the most degraded area. This finding coincides with previous studies (Ortiz *et al.,* 2020), and could be associated with the history of agricultural burning in this area, mainly of potato cultivation (*Solanum tuberosum* L?), which generates pyrogenic carbon. This form of carbon is evidenced by the presence of charcoal fragments and an intense black color in the soil samples. Pyrogenic carbon is highly resistant to oxidation due to its polyaromatic structure, being a persistent fraction of the total carbon, making it an important long-term carbon reservoir (Lavalle *et al.,* 2019b).

Furthermore, silvopastoral management conditions showed a higher percentage of carbon compared to the non-silvopastoral treatment (Op > SC > SOp > Ctr). This suggests that silvopastoral systems promote soil recovery, increasing carbon accumulation, possibly through the increase in the amount of available organic matter contributed by animals to the system. In general, the estimated carbon concentrations were higher than those observed in other soil conservation systems, an example of this is the study by Poblete- Grant *et al*. (2020), which reports organic carbon levels in Andisol soils under grasslands in southern Chile with prolonged application of poultry manure, which were lower than those found in this study. Our results are, however, similar to those reported by Gómez *et al*. (2020), who evaluated these parameters under similar agroforestry conditions in a temperate native forest in Argentina, working with the tree species *Nothofagus antartica*. This species, like *N. obliqua*, is a deciduous tree that contributes a continuous level of leaf litter to the soil throughout the year, which is related to high carbon levels inputs.

The total N% showed significant differences between the canopy opening conditions evaluated (ANOVA F = 19.90; df = 3.32; p = 0.0001) and soil depths (ANOVA F = 25.17; df = 3.32; p = 0.0001) (Figure 5b). The total N% of (0-10cm) varied between 0.45 and 1.0 (±0.15) where Op˃ SC˃ SOp˃ Ctr. The values obtained in this study are higher than those reported by Crovo *et al*. (2021) for native forest Andisols. However, the inorganic forms of nitrogen available to plants NO3 and NH4, were lower than the minimum requirements necessary for plant growth (Table S2). Average NO₃⁻ values ranged from 7.77 to 37.73 mg/kg (Ctr > SC > Op > SOp), while NH₄⁺ values ranged from 3.01 to 12.83 mg/kg (SOp > SC > Op > Ctr), respectively. Previous studies have shown a similar trend or behaviour at the same site in previous years (Alfaro et al., 2018; Ortiz et al., 2020), although an increase in NH₄⁺ levels as a form of bioavailable nitrogen would have been expected due to the incorporation of animal feces.

In terms of C sequestration, the results indicate that the Op treatment is the most effective, as it presents the highest SOC stocks at all depths (Figure 6). The highest Op levels with respect to the rest of the treatments were observed in the surface horizons 0-10 cm and deep horizons 30-60 cm, where values of 84.2 Mg C ha⁻¹ and 189.6 Mg C ha⁻¹ were reached, respectively. On the other hand, the SOp and SC treatments also showed high values at all depths analysed, although not as high as Op, this could be due to differences in management practices associated with agricultural burning, which influence the dynamics of carbon in the soil. The Ctr presented the lowest C stock values with respect to the rest of the treatments, especially at the depth of 30-60 cm, where it barely reached values close to 90 Mg C ha⁻¹. This suggests that soil in unmanaged degraded native forests has a lower capacity to retain carbon, and that the implementation of practices such as SPS are effective in promoting long-term C storage.

**Figure 6.**
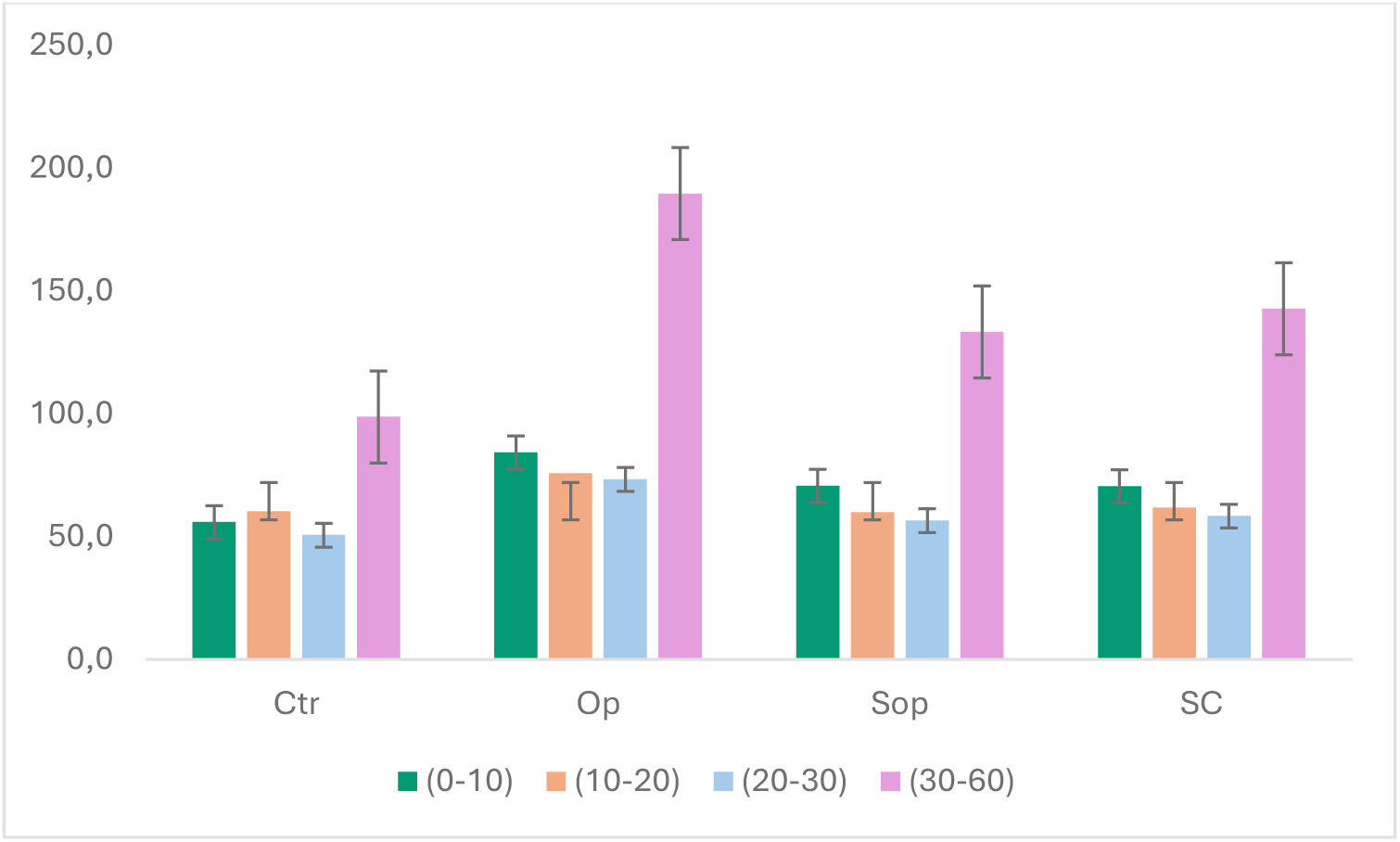
SOC stocks (Mg ha^-1^) at depths 0-10, 10-20, 20-30 and 30-60 cm for Ctr, Op, SOp and SC respectively.

The results revealed that the C/N ratio of the unfractionated samples did not show significant differences in terms of canopy openness (ANOVA F = 1.93; df = 3.32; p = 0.1451) but did differ in terms of soil depth (ANOVA F = 6.37; df = 3.32; p = 0.0016) (Figure 7a). The C/N ratio of the unfractionated samples varied between 12.66 and 19.25 with an average of 15.6 ±1 which is within the distribution of C/N averages (9.9 to 25.8) found for soils around the world (Batjes, 2014), although it is below the C/N ratio reported by Katsumi *et al*. (2015) who worked with Andisols (n = 14) under different land uses and found an average C/N ratio of 26.3. These results for the site could be highly related to the quality of the substrate, since the tree species present at the site is deciduous and its litter has a relatively lower lignin content (lignin:N ratio of 21.79 and, consequently, lower C:N ratios (Decker and Boerner 2003).

**Figure 7:**
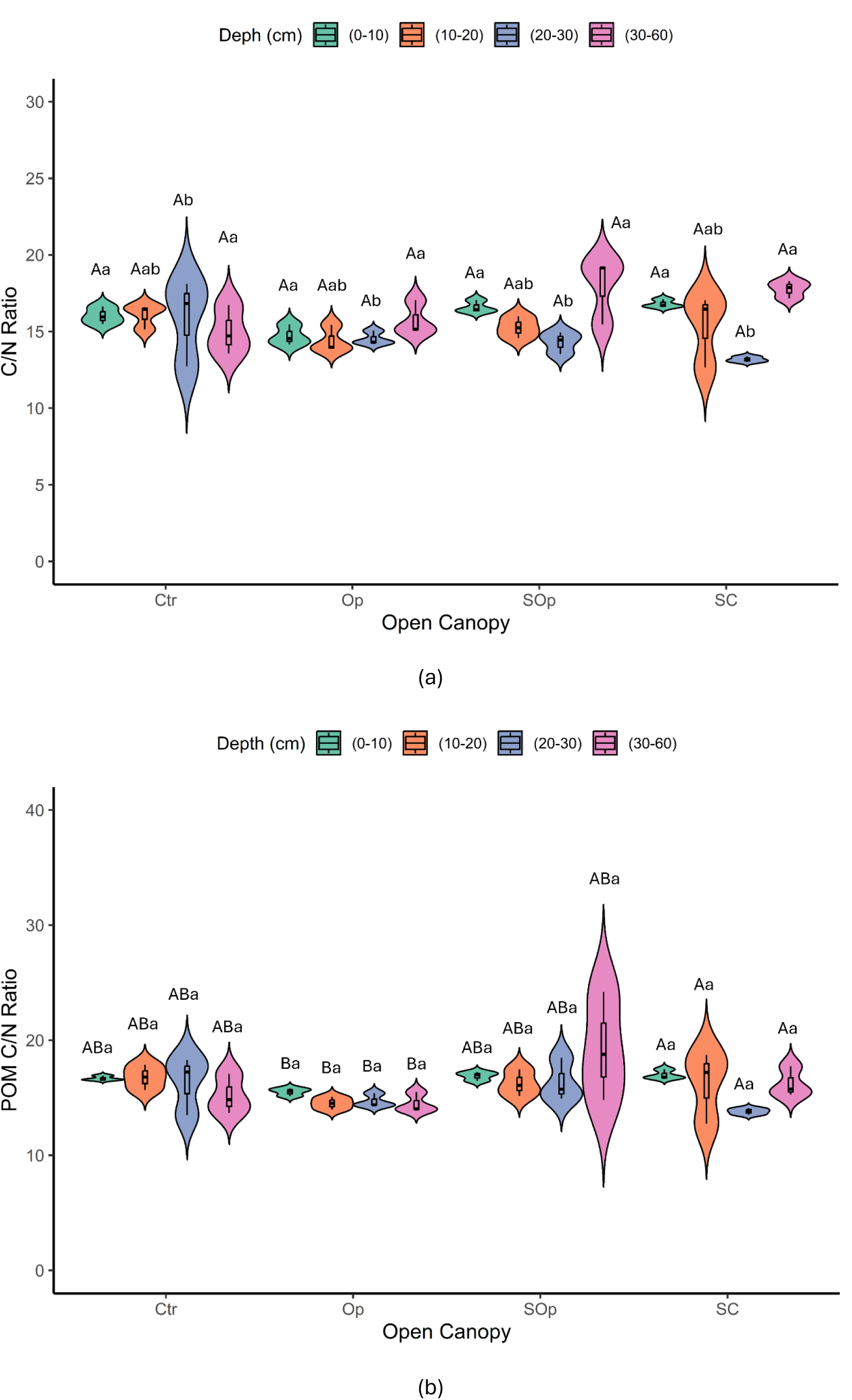
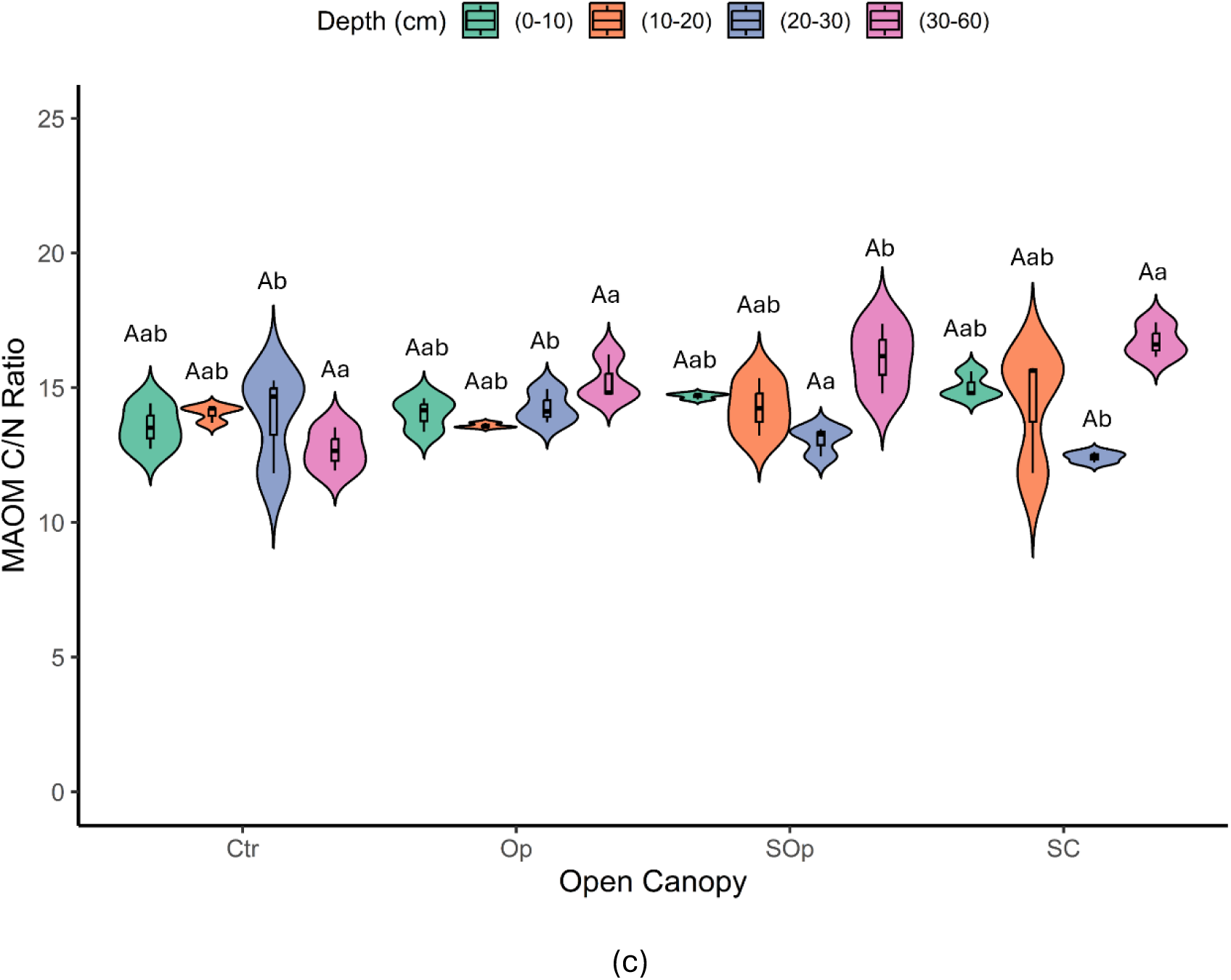
Violin plot of the variables (a) C/N ratio (b) POM C/N ratio and (c) MAOM C/N ratio as a function of canopy openness and soil depth factors. Bars represent the meaning of each variable, and whiskers indicate the standard error of the mean. Different capital letters mean significant differences between conditions, while lowercase letters indicate significant differences between depths. Analysis of variance (ANOVA) was used to determine significant differences. Differences were considered significant at a p < 0.05 level.

The C/N ratio of the POM fraction showed significant differences depending on the canopy opening conditions (ANOVA F = 3.68; df = 3.32; p = 0.0220) (Figure 7b). On the other hand, the C/N ratio of the MAOM fraction showed significant differences both with depth (ANOVA F = 6.93; df = 3.32; p = 0.0426) and with canopy opening (ANOVA F = 3.05; df = 3.32; p = 0.0010) (Figure 6c). In the particulate fraction, the highest C/N ratios were observed at depths (30-60 cm), particularly in SC, while in the surface horizons (0-10 cm), the values were more uniform between treatments. On the other hand, in the mineral fraction, C/N values were more homogeneous throughout the depths, although the surface horizons showed lower C/N ratios, with Ctr being the treatment with the lowest values.

Overall, the C/N ratio of MAOM (11-17, mean = 14.24 ± 1) was very similar to that of POM (12-24, mean = 16.02 ± 2), although more contrasting values between both fractions would have been expected (Figure 7c). As reported by Lavallee *et al*. (2020), POM generally presents C/N ratios between 10 and 40, while MAOM has narrower C/N ratios, between 8 and 13. This would be expected, given that the C/N ratios of POM and MAOM largely reflect the characteristics of the source material: POM is associated with the C/N of plant litter, while MAOM corresponds to the C/N of microbial necromass. Plant litter shows high variability in its C/N ratio (e.g., from 40 to 120 in a subset of NEON sites; Hall *et al.,* 2020), while soil microorganisms have a significantly lower and restricted C/N ratio (between 3 and 15; Strickland and Rousk, 2010). This analysis would be based on the hypothesis that MAOM is dominated by microbially derived organic matter, however, POM and MAOM fractions are also known to contain mixtures of faster and slower cycling C pools with likely contributions from both plant and microbial detritus (Hall *et al*., 2015; Poeplau et al., 2018; Angst *et al*., 2021). A study by Wenjuan *et al*. (2022) suggests that plant-derived organic matter may also contribute substantially to MAOM, especially in humid forests (with MAOM C/N > 15), which could explain the similarity observed between fractions in this study.

Cnox concentrations in unfractionated samples varied significantly with respect to opening condition (ANOVA F = 19.90; df = 3.32; p = 0.0001) and depth (ANOVA F = 19.90; df = 3.32; p = 0.0001) (Figure 8a). Total oxidizable carbon decreases with depth in all treatments. The highest values are found in the Op and SC treatments in the surface horizons despite being the most different treatments in terms of degradation condition, but this is not the case at the depth of 30-60 cm where the differences between all treatments tend to be less marked.

**Figure 8:**
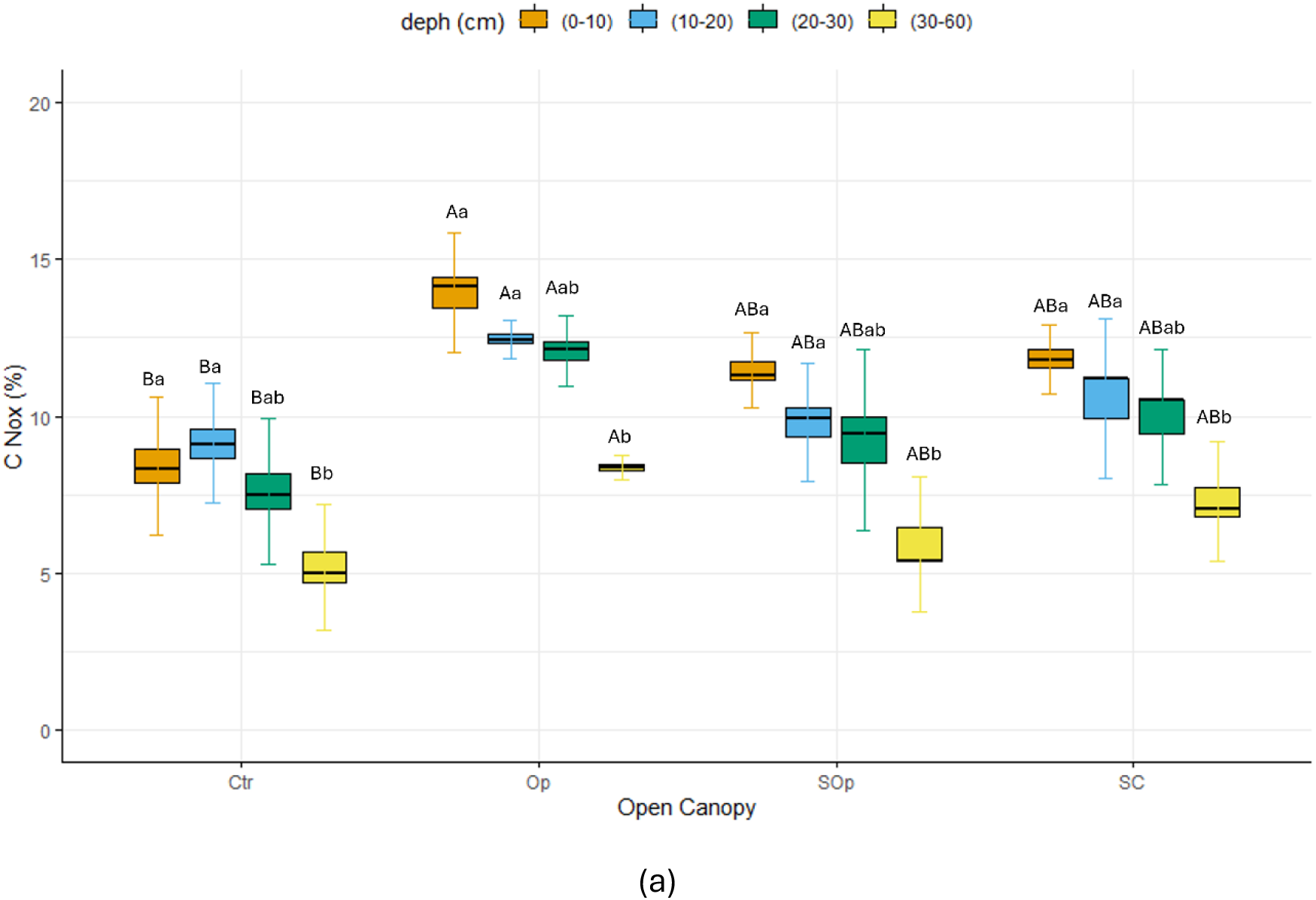
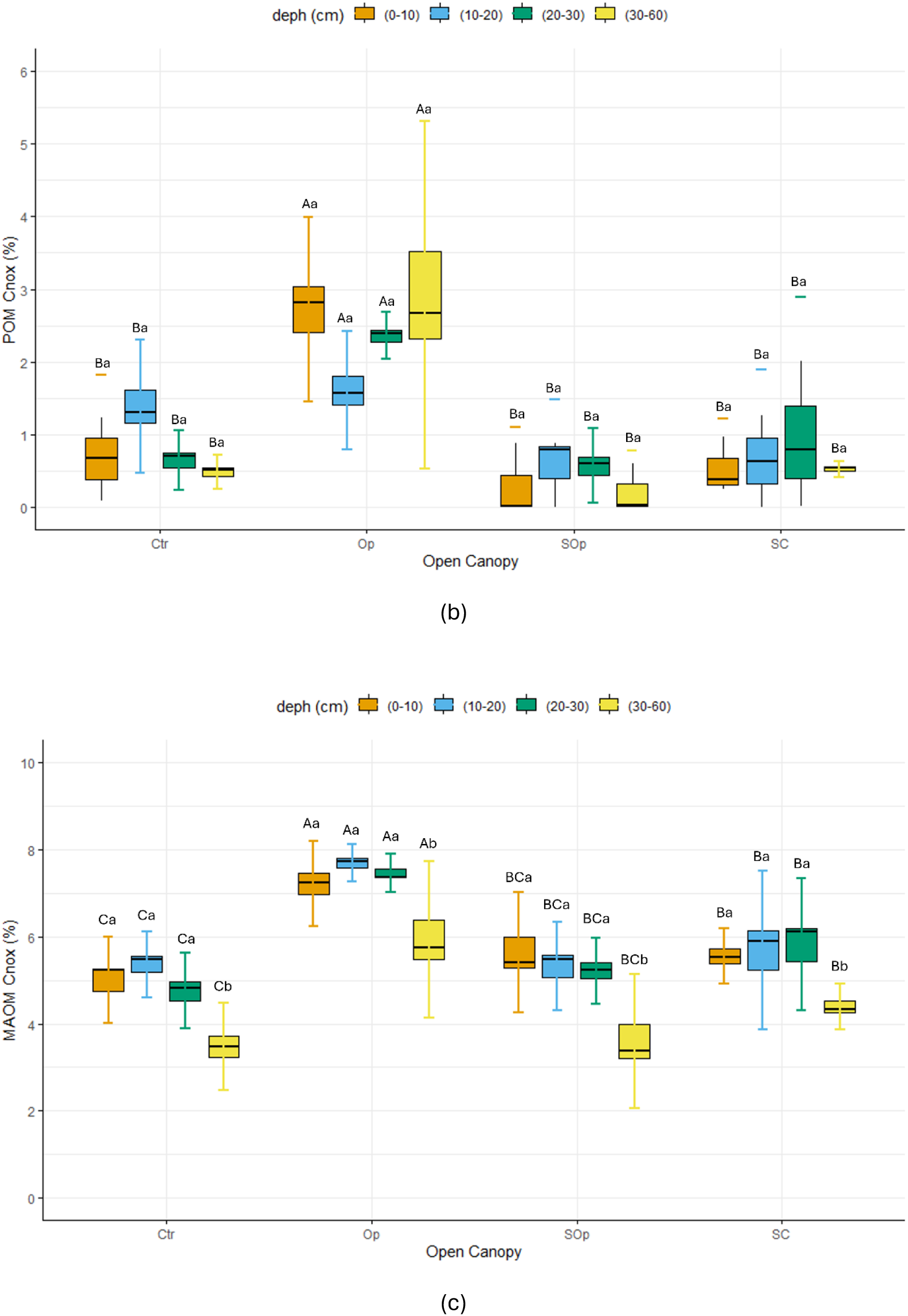
(a) Non-oxidizable carbon (b) POM non-oxidizable carbon and (c) MAOM non-oxidizable carbon as a function of canopy openness and soil depth factors. Bars represent the mean of each variable, and whiskers indicate the standard error of the mean. Capital letters mean significant differences between conditions, while lowercase letters indicate significant differences between depths. Analysis of variance (ANOVA) was used to determine significant differences. Differences were considered significant at p < 0.05.

The concentration of POM Cnox and MAOM Cnox showed significant differences depending on the canopy opening conditions (ANOVA F = 32.33; df = 3,32; p = 0.0001 and ANOVA F = 42.36; df = 3,32; p = 0.0001, respectively). In addition, the MAOM Cnox fraction also showed significant differences depending on the depth (ANOVA F = 19.27; df = 3,32; p = 0.0001) (Figure 8b and 8c). In general, the values of POM Cnox and MAOM Cnox tend to be higher in the surface horizons (0-10 cm) and progressively decrease with depth, following a pattern similar to that observed in the unfractionated samples.

As reported by Sierra *et al*. (2013), it is expected that as the soil profile goes deeper, the fraction of non-oxidizable carbon (Cnox) will increase proportionally, reflecting the accumulation of more recalcitrant and stable forms of organic matter in the deeper layers. This phenomenon is due to the presence of recalcitrant compounds such as lignin, tannins (Kraus et al., 2003), cutins and suberins (Tegelaar *et al.,* 1989), which are more abundant in the roots (Goering and Van Soest, 1970). On the other hand, it is expected that the concentrations of oxidizable carbon (Cox) will be higher in both fractions (POM and MAOM) in the surface horizons, due to the contribution of organic carbon from leaf litter.

In this study, both Cox and Cnox showed the highest concentrations in the surface horizon (0-10 cm) (Figure S1a). Wenjuan *et al*. (2022), in a study that included a large transect of soil types, found that forest soils tend to vary the chemical and isotopic properties of the organic matter fractions of the soil and that it occurs mainly in the surface horizons. These authors found that in the surface horizons of forest soils there is a high abundance of aliphatic compounds (aliphatic C–H/C=O) compared to carbonyl groups (C=O), which are associated with more oxidized compounds. These observations reinforce the relationship between carbon dynamics in the different organic matter fractions and the fact that the C stabilization processes in our study are not only occurring at depth.

In terms of canopy opening, the Op treatment showed the highest concentrations of both fractions. However, MAOM Cnox concentrations are more homogeneous throughout the depths compared to the POM fraction, which presents greater variability. This behavior could be related to the different dynamics of carbon stabilization in each fraction, where MAOM is more associated with stabilized microbial organic matter, while POM reflects a more direct influence of litter and other materials in initial decomposition processes. In addition, Cnox is determined as a more stable fraction of C and is expected to include highly recalcitrant forms of C, including pyrogenic C or black carbon, and/or organic compounds strongly associated with the clay fraction. This is why the Op condition, despite being the most degraded, presents the highest values and this is associated with the historical anthropogenic practices of burning stubble.

PCA explained 77.1% of the total variation in the data with the first two components (Figure 9). The results revealed clear clusters for biological and C and N analyses in the samples and their fractions indicating that agroforestry practices have a significant impact on SOC properties and dynamics.

**Figure 9:**
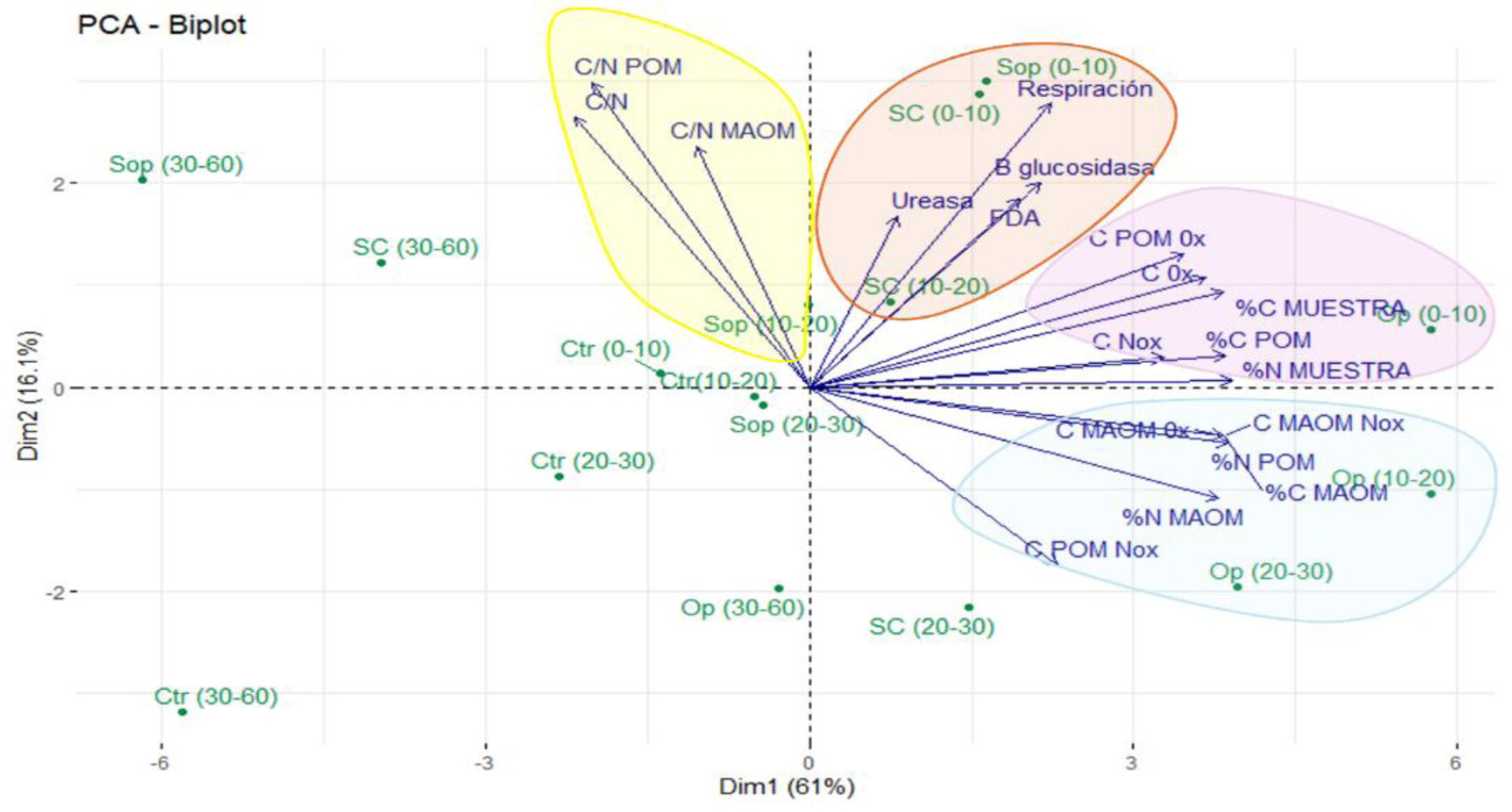
Principal component analysis (PCA) of soil chemical and biological indicators under different canopy openness conditions (Ctr, Op, SOp, and SC) and soil depths (0-10, 10-20, 20-30, and 30-60 cm). Arrows represent the original variables projected into the space of the first two principal components (Dim1 and Dim2), which explain 77.1% of the total variance, respectively. Colored ellipses group observations according to their similarities, providing information on the relationship between variables under different canopy openness conditions and depths. Arrows indicate the contribution of each response variable to the variability explained by the principal components, highlighting the differences in parameters under each treatment.

At shallow depths of 0-10 cm and 10-20 cm for the SOp and SC systems, they are grouped together, showing an association with the variables of enzymatic activity and soil respiration. This indicates that silvopastoral systems at shallow depths present a high biological activity, which is consistent with their greater exposure to the input of organic matter and conditions that favor microbial activity. The Op system and the depths of 20-30 cm and 30-60 cm tend to group together, showing an association with the content of oxidizable and non-oxidizable carbon in different fractions. This suggests that in this system, although it is more degraded, the accumulation of stable carbon could be promoted by historical management practices (such as burning), which generate pyrogenic carbon resistant to degradation. The Ctr is not grouped with the variables of carbon content and enzymatic activity. This suggests a lower level of carbon and lower biological activity, which could be determined since this system does not present silvopastoralism.

## 4. Conclusions

This study highlights the critical role agroforestry systems based on native forests can play in restoring degraded soils. The increased levels of stable carbon and nitrogen observed in silvopastoral system (SPS) treatments, compared to the control (Ctr), emphasize the potential of these practices to enhance long-term carbon sequestration and improve soil quality in degraded landscapes. The distinct dynamics of carbon across various organic matter fractions, coupled with the observation that carbon stabilization in this study is not confined to deeper soil layers, may be characteristic of humid forest ecosystems. Future research should incorporate more advanced methodologies, such as fractionation techniques that separate soil organic matter (SOM) by particle size or density, and targeted analyses to identify pyrogenic carbon. These approaches would enable a more nuanced understanding of the mechanisms and pathways driving SOM formation and stabilization, thereby advancing our knowledge of its dynamics across diverse ecological and management contexts.

